# A revised calcium-dependent model of theta-burst transcranial magnetic stimulation

**DOI:** 10.1101/2022.04.15.488478

**Authors:** K. Ma, J. C. Rothwell, S. M. Goetz

## Abstract

Calcium dependency is presently an essential assumption in modelling the neuromodulatory effects of transcranial magnetic stimulation (TMS). Among the various neuromodulatory TMS protocols, theta-burst stimulation (TBS) at present is the fastest intervention to generate strong effects. A decade ago, Y.Z. Huang et al. developed a first neuromodulation model to explain the bidirectional effects of TBS based on postsynaptic intracellular calcium concentration elevation. We discover, however, that the published computer code is not consistent with the model formulations in the corresponding paper. Further analysis confirms that the computer model with an index confusion was used for fitting the experimental results, running the simulation, and plotting the corresponding figures in the original publication. This paper intends to fix the computer code and additionally create a non-convex optimisation solution for re-calibrating the model. After re-calibration, the revised model outperforms the initial model in accuracy describing the MEP amplitudes of TBS-induced after-effects under specific situations.

## 1. Introduction

Repetitive transcranial magnetic stimulation (rTMS) is a non-invasive brain stimulation technique for modulating neural circuits in the human cortex (Barker et al., 1985). It is widely used in experimental brain research and clinical applications, such as the therapy of major depression, obsessive compulsive disorder, addiction, migraine, stroke, and epilepsy as well as diagnosis (Wagner et al., 2007; Mantovani et al., 2010; Eldaief et al., 2013; Luan et al., 2014; Rossini et al., 2015; Valero-Cabré et al., 2017; Beynel et al., 2019). To date, a variety of rTMS protocols can induce neuromodulatory effects, such as facilitation and inhibition of neural activity (Di Lazzaro et al., 2011). Particularly, theta-burst stimulation (TBS), a patterned form of rTMS introduced by Huang et al., was enabled by the advancement of stimulator technology that is capable to generate high-frequency trains of stimuli (Huang et al., 2005; Goetz, Deng, 2017). TBS protocols entail major advantages over traditional rTMS protocols generating strong long-lasting effects after substantially shorter interventions and lower stimulation pulse amplitudes (Suppa et al., 2016a; Chung et al., 2015; Wischnewski, Schutter, 2015; Turi et al., 2021). After a successful clinical trial, it was recently approved for the treatment of major depression (Blumberger et al., 2018). Intensive therapy with multiple sessions a day may further increase efficacy and accelerate treatment (Cole et al., 2020).

TBS protocols typically share the same fundamental building block that consists of 5 Hz bursts of three pulses repeated at 50 Hz and differ in the inter-train intervals and the number of bursts and trains (Figure 1). Two TBS protocols dominate: continuous TBS (cTBS) and intermittent TBS (iTBS). So-called intermediate TBS (imTBS) is not frequently investigated because it has little to no detectable effect if administered over the primary motor cortex (Huang et al., 2005). Most quantitative studies use the primary motor cortex as a model circuit, where motor-evoked potentials (MEP) in response to constant single-pulse TMS can demonstrate changes of excitability. 600 pulses of iTBS (iTBS600) on average increase MEPs, while the same number of cTBS pulses (cTBS600) decreases MEPs (Suppa et al., 2016b; Corp et al., 2020). Moreover, numerous results from human studies also demonstrate that voluntary muscle contraction before, during, and after the TBS intervention can interact with the after-effects of TBS on MEPs (Huang et al., 2005; Gentner et al., 2008; Gamboa et al., 2011; Haeckert et al., 2021). Huang et al. (2011) proposed a compact mathematical model of TBS-induced after-effects based on the assumption of glutamatergic dynamics of synaptic plasticity. The Huang model already takes into account protocol-dependent and contraction-dependent factors at the system level. With the calibration reported by Huang et al. (2011), it achieved to successfully describe the time courses and polarities of the after-effects of cTBS300/600 and iTBS600 with respect to conditions of prior or post voluntary contractions.

**Figure 1:**
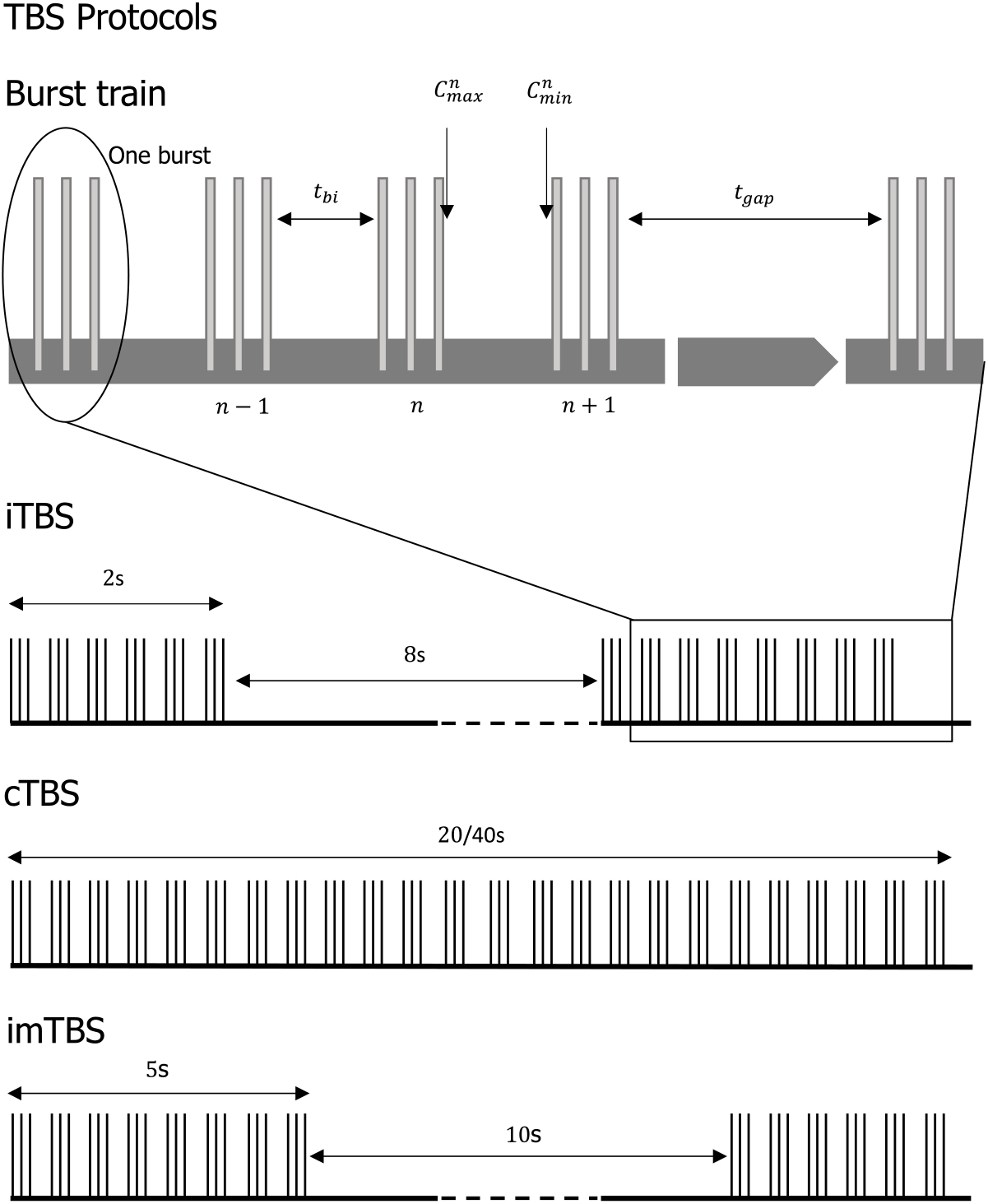
Schematic diagram of theta-burst protocols. Each train consists of 5 Hz bursts with three pulses repeated at 50 Hz. Bursts are equally spaced with an interval of *t*_bi_ between two bursts. The maximum postsynaptic cal-cium level, 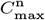, occurs at the time right after the *n*-th burst, while the minimum calcium level, 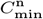, occurs at the time right before the (*n* + 1)-th burst. Trains are equally spaced with an interval of *t*_gap_ between two trains. Each protocol contains *T* trains, each of which contains *B*_t_ bursts. For example, iTBS is represented by *B*_t_ = 10, *T* = 20, *t*_bi_ = 0.16 s and *t*_gap_ = 8 s; cTBS by *B*_t_ = 100 *or* 200, *T* = 1, *t*_bi_ = 0.16 s and *t*_gap_ = 0 s; imTBS uses *B*_t_ = 25, *T* = 8, *t*_bi_ = 0.16 s and *t*_gap_ = 10 s.

However, an in-depth analysis discovered that the published computer code^1^ is not consistent with the model equations in the paper. The paper calculates the rate of increase of the calcium level at the current state for the facilitatory *substance* by subtracting the calcium level at the previous burst from that at the current burst in Huang et al. (2011). The computer code, however, subtracts the calcium level at the one before the previous burst from that at the current burst, skipping one burst in the middle. Thus, the index usage in the article and in the computer code used for the calibration in the earlier article appear to show discrepancies. Further analysis confirms that the computer model with the index confusion was used for the calibration to the experimental data. Consequently, the graphs in the paper is based on the code rather than the correct equations presented in the paper (Huang et al., 2011). Fixing this error will result in other difference equations in the code and will require new calibration.

Even though the initial computer code might still reproduce experimental observations to some degree—which would be important to retain a rich body of research derived from the model— the accuracy and the scope of the model may be affected. Furthermore, parameters of the model as well as the internal variables cannot be interpreted correctly anymore. Therefore, the purpose of this paper is to correct, modify, and re-calibrate Huang model to the experimental data, while also adding clearer explanations and a specific statement on the model’s validity.

The further text will introduce a detailed procedure of construction of the model, including key assumptions. Subsequently, we will compare the results of the revised model with the initial one and discuss the differences. The article will conclude with the limitations of this model as well as a brief guideline of the future considerations.

## 2. Structure of Huang’s Model

### 2.1. Fundamental Design Aspects

Although TBS may affect a variety of neurotransmitters (Huang et al., 2007; Stagg et al., 2009; Li et al., 2019), the Huang model concentrates on glutamatergic synapses and N-methyl-D-aspartate (NMDA) receptors in particular by modelling the postsynaptic intracellular calcium concentration ([Ca^2+^]_*i*_) and corresponding downstream protein kinases/phosphatases (Yang et al., 1999; Lee et al., 2000; Huang et al., 2011). It assumes that TBS-induced postsynaptic Ca^2+^ influx can simultaneously activate both facilitatory and inhibitory mechanisms, quantified through *substances*, the sum of which determines the polarity and magnitude of the after-effects on MEPs (Huang et al., 2011). The facilitatory *substance* depends on the increase rate of the postsynaptic calcium concentration, while the inhibitory *substance* depends on sustained quantities of Ca^2+^ entry (Yang et al., 1999). As a result, the model of TBS-induced after-effects is divided into three stages that mimic the biological responses in synapses. All TBS protocols in this model are applied at a stimulation pulse strength of 80% active motor threshold (AMT) or 70% resting motor threshold (RMT) (Huang et al., 2005, 2008, 2011; Gentner et al., 2008).

### 2.2. Stage I: Changes of calcium level due to bursts

The model assumes that each burst induces postsynaptic Ca^2+^ influx of the small amount of *C*_b_ (arbitrary unit). The level of Ca^2+^ decays exponentially after each burst. Hence, the level of Ca^2+^ at a time point *t* after a single burst will be

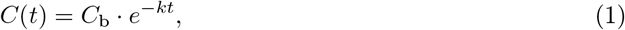

where *k* is the decay time constant for Ca^2+^.

The following supposes that the level of calcium does hardly decay during a burst due to its short duration. As shown in Figure 1, the maximum level of Ca^2+^ is

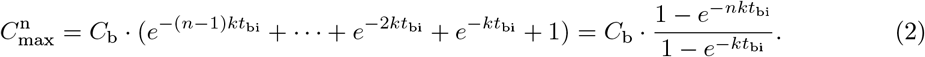

The minimum level of Ca^2+^ (before the *n* + 1-th burst) is defined as

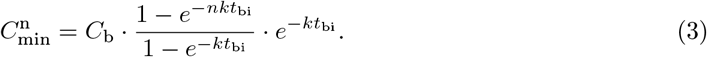

where *t*_bi_ is the time between two bursts. In this paper, we selected 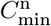 to represent the calcium state after each burst.

### 2.3. Stage II: Accumulation of facilitatory and inhibitory substances

Both *substances* decay exponentially with time. The facilitatory *substance* is defined as the accumulation of the rate of increase in the postsynaptic calcium level. Thus, the amount of facilitation after *n* regular bursts follows

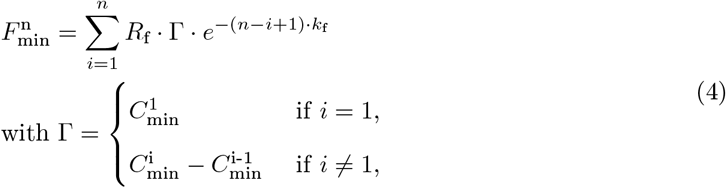

where *R*_f_ is a proportionality constant of facilitation and *k*_f_ its decay time constant.

The inhibitory *substance* is defined as the accumulation of the calcium level. Hence, the amount of it after *n* regular bursts is calculated as

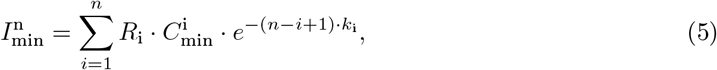

where *R*_i_ is a proportionality constant of inhibition and *k*_i_ its decay time constant.

### 2.4. Stage III: Facilitatory and inhibitory after-effects on MEPs

Stage III describes the relationship between the after-effects on MEPs and time *t* after TBS via a nonlinear sigmoidal function formulated as

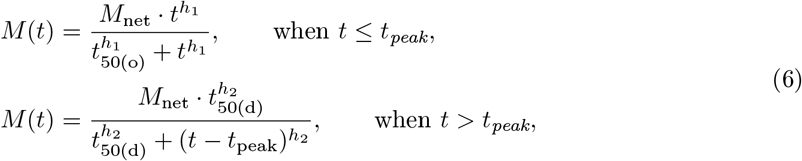

where *h*_1_ and *h*_2_ are power coefficients that describe the steepness of the sigmoid curves, and *M*_net_ is the net effect. In this model, we have *t*_50(o)_ = *k*_1_ · *t*_peak_, *t*_50(d)_ = *k*_2_ · *t*_peak_ in which *k*_1_ ∈ (0, 1) and *k*_2_ ∈ (1, ∞) because of *t*_50(o)_ *< t*_peak_ and *t*_50(d)_ *> t*_peak_.

The peak time for the inhibitory after-effect defined in the initial model is

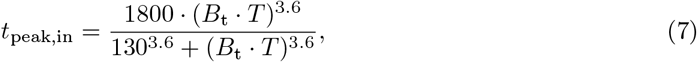

where *B*_t_ is the number of bursts in each train and *T* is the number of trains delivered in one TBS session (Huang et al., 2011). The peak time for the facilitatory after-effect, *t*_peak,fa_, is approximately one fifth of *t*_peak,in_ according to previous observations (Huang et al., 2011).

Suppose that there are *T >* 1 regular trains and each train contains *B*_t_ *>* 1 regular bursts (see Figure 1), the *substances* of facilitation 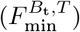 and inhibition 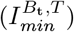 right after *T* trains are calculated per

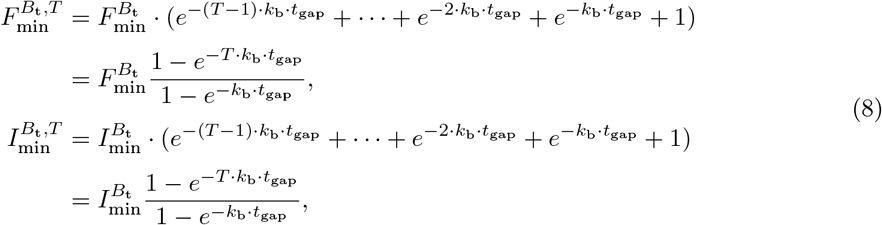

where *k*_b_ is the decay constant for the decline between trains and *t*_gap_ is the inter-train interval. Hence, the net effect is 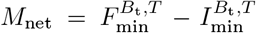 (Huang et al., 2011). Furthermore, the after-effect *M* (*t*) has two different sets of parameters, 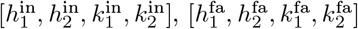, respectively for modelling after-effects of inhibition and facilitation.

### 2.5. Metaplastic effects of voluntary contraction

Abraham, Bear (1996) suggested that the subsequent synaptic plasticity can be modulated by prior synaptic activity, which is termed metaplasticity. Prior voluntary contraction—as also used for AMT titration—may modify synaptic functionality to cause metaplasticity (Abraham, 2008; Citri, Malenka, 2008; Wankerl et al., 2010). The model assumes that a prior contraction lowers the value of *C*_b_, that the proportionality (*R*_i_) and decay constant (*k*_i_) of the inhibitory *substance* 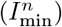 are inversely proportional to 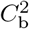, and that the decay constant (*k*_f_) of the facilitatory *substance*) 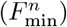 is inversely proportional to *C*_b_ (Huang et al., 2011).

Moreover, due to the experimentally observed elimination of inhibition after voluntary contraction after TBS (Huang et al., 2008), Huang et al. postulated that the inhibitory after-effect is blocked then, while the facilitatory after-effect is preserved, i.e., 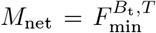 (Huang et al.,2011). Note that the inhibitory after-effect curve is used only when there is no prior contraction or 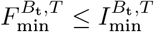 while the facilitatory after-effect curve is used for other conditions (Huang et al., 2011).

## 3. Methodology for Parameter Calibration

We collected the experimental data of mean value of MEPs from papers by digitizing plots from studies listed in Table 1 (Huang et al., 2005, 2008; Gentner et al., 2008).^2^ As suggested by Huang et al. (2011), all measurements of normalised MEPs are shifted down by 1 (to delete the baseline) and scaled up tenfold. The time is in seconds.

**Table 1:** Overview of TBS experiments that have been used for re-calibration in the present article. (a) AMT: active motor threshold; (b) RMT: resting motor threshold; (c) Data code is used to assign plots to corresponding data in following sections; (d) Sample size: the number of healthy subjects.

| Study | TBS protocol | Sample Size | TBS intensity | Contraction condition | After-effect on MEP | Data Code |
| --- | --- | --- | --- | --- | --- | --- |
| Huang et al. (2005) | cTBS300 | N=9 | 80% AMT | Prior contraction | Inhibition | HU05 |
|  | cTBS600 | N=9 |  | Prior contraction | Inhibition |  |
|  | imTBS600 | N=9 |  | Prior contraction | None |  |
|  | iTBS600 | N=9 |  | Prior contraction | Facilitation |  |
| Huang et al. (2008) | cTBS300 | N=9 | 80% AMT | Prior and post contraction | Facilitation | HU08 |
|  | iTBS600 | N=7 |  | Prior and post contraction | Facilitation |  |
| Gentner et al. (2008) | cTBS300 | N=16 | 70% RMT | None | Facilitation | GE08 |
|  | cTBS600 | N=9 |  | None | Inhibition |  |

While the computer codes of Stages I and III are not modified, we re-calibrate the parameters at Stage II of the revised model, which exactly matches the equations in the printed article of Huang et al. (2011), through nonlinear least-squares parameter estimation with ℒ^2^ penalty. We define that *R*_f_ = *D*_1_, *K*_f_ = *D*_2_/*C*_b_, 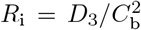, and 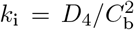. Due to the dependency on voluntary muscle contraction (Gentner et al., 2008; Huang et al., 2008), we have two factors: (a) with/without prior contraction (*Contr*_*prior*_ = 1 *or* 0); (b) with/without immediate post contraction (*Contr*_*post*_ = 1 *or* 0). Therefore, the net effect *M*_net_ for a TBS protocol with muscle conditions is a function *M*_net_(**Pro, X**), thereby defining the static after-effect model at time *t* after stimulation as 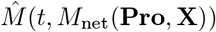, where **Pro** = [*B*_t_, *T, t*_bi_, *t*_gap_, *Contr*_*prior*_, *Contr*_*post*_] and **X** = [*D*_1_, *D*_2_, *D*_3_, *D*_4_].

Suppose that we have in total *N* measurements of the after-effects of different TBS protocols, the *j*-th measurement *A*^*j*^, for a particular TBS protocol **Pro**^*j*^, was measured at the *j*-th time point *t*^*j*^. Hence, the sum of the squares of the residuals between the measurement *A*^*j*^ and the curve-fit model 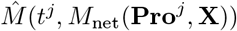 with regularization follows

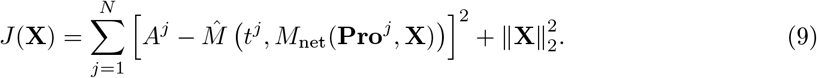

Considering the constraints, all parameters should be real positive numbers. In conclusion, we formulate a non-convex nonlinear programme for optimally calibrating the parameters of the revised Huang model as

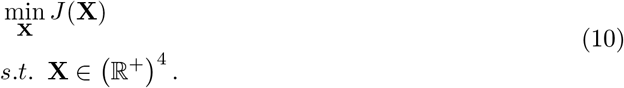

We used the Levenberg–Marquardt algorithm as implemented in Matlab (v2021b, The Mathworks, USA) for solving Eqs. (9) and (10). Due to the nonlinearity of this cost function, the global optimality is not guaranteed. The initial and re-calibrated parameters are listed in Table 2. (Please see Appendix Appendix A for the code written in MATLAB.)

**Table 2:** Summary of initial parameters in Huang’s initial model and new parameters after optimisation in the revised Huang model (Huang et al., 2011).

| Parameter | Initial | Re-calibrated | Biological relevance |
| --- | --- | --- | --- |
| **Parameters for Stage I** |  |  |  |
| $C_b(0)$ | 3 | 3 | $\text{Ca}^{2+}$ influx after a burst without prior contraction. |
| $C_b(1)$ | 1 | 1 | $\text{Ca}^{2+}$ influx after a burst with prior contraction. |
| $k$ | 1.2 | 1.2 | Decay constant of $\text{Ca}^{2+}$ level. |
| **Parameters for Stage II** |  |  |  |
| $R_f$ | 1 | 1.45 | Proportionality constant of facilitation. |
| $k_f$ | $0.01/C_b$ | $0.005/C_b$ | Decay constant of facilitation. |
| $R_i$ | $0.14/C_b^2$ | $0.123/C_b^2$ | Proportionality constant of inhibition. |
| $k_i$ | $0.055/C_b^2$ | $0.07/C_b^2$ | Decay constant of inhibition. |
| $k_b$ | 0.1 | 0.1 | Decay constant for the declines during intertrain intervals. |
| **Parameters for Stage III** |  |  |  |
| $h_1^{\text{in}}/h_1^{\text{fa}}$ | 2.5 / 3 | 2.5 / 3 | Power coefficients of the sigmoidal curve $M(t)$, describing the steepness of curve slopes. |
| $h_2^{\text{in}}/h_2^{\text{fa}}$ | 4 / 2 | 4 / 2 | |
| $k_1^{\text{in}}/k_1^{\text{fa}}$ | 0.2 / 0.25 | 0.2 / 0.25 | Proportionality constants of $t_{\text{peak}}$. |
| $k_2^{\text{in}}/k_2^{\text{fa}}$ | 1.1 / 2 | 1.1 / 2 | |

## 4. Results

This section looks at the root-mean-square error (*RMSE*) and the pseudo R-squared value to quantitatively evaluate the goodness-of-fit for the initial and revised models, which are calculated as

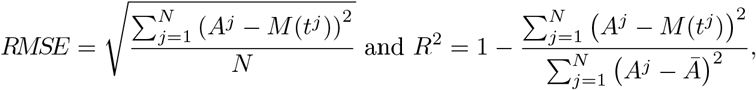

where *Ā* is the mean of the measurements (Efron, 1978). Moreover, we use the standard deviation to represent the distribution of residuals, specifically

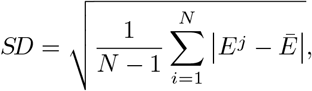

where *E*^*j*^ = *A*^*j*^ − *M* (*t*^*j*^) and *Ē* is the mean of the residuals.

The pseudo R-squared of the initial model 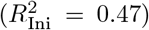 is lower than of the revised model 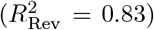, indicating that the updated model and parameters at Stage II reduce the unaccounted variability and improve the quantitative description of the after-effects (Figure 2). Consistently, the revised Huang model has a lower RMSE value (*RMSE*_Rev_ = 1.81) than the initial one (*RMSE*_Ini_ = 3.18) indicating a better fit in terms of absolute errors. Moreover, Figure 3 illustrates that the initial model has a wider error span [−8, 9] compared to the relative small error span [−5, 6] of the revised Huang model, and their standard deviations are respectively *SD*_Ini_ = 2.99 and *SD*_Rev_ = 1.81. In conclusion, the revised model has an advantage over the initial model in following the data without any additional degrees of freedom.

**Figure 2:**
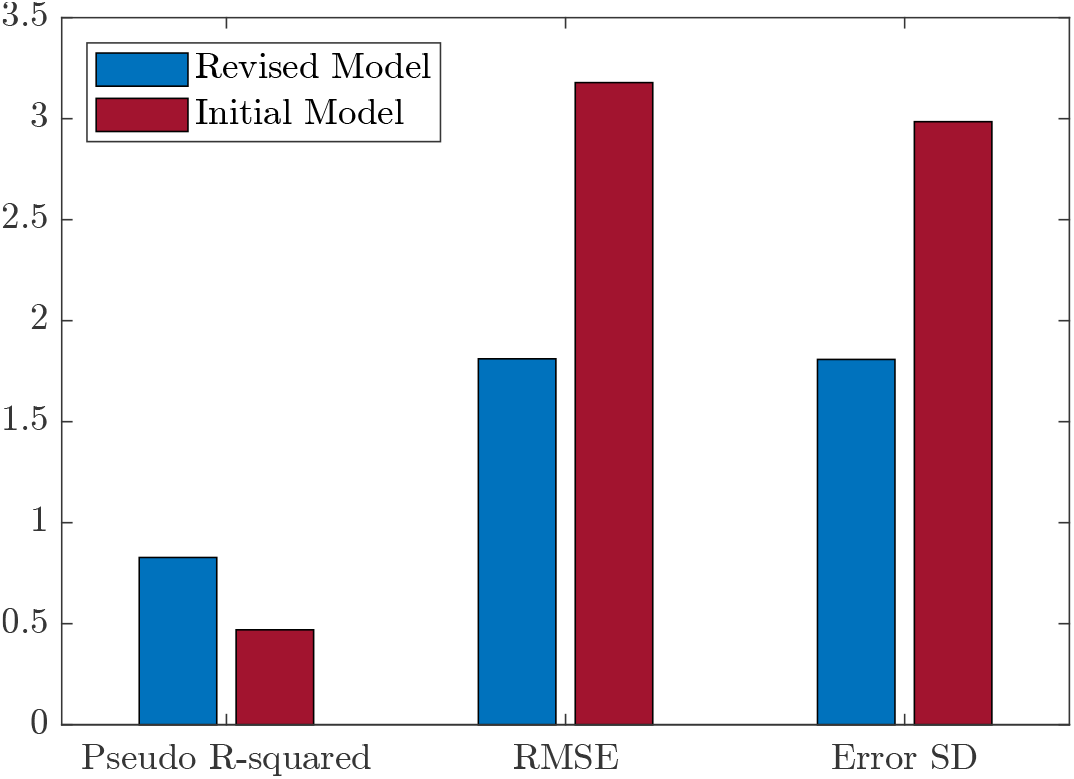
Evaluation of the goodness-of-fit of initial and revised models. RMSE: root-mean-square error; Error SD: standard deviation.

**Figure 3:**
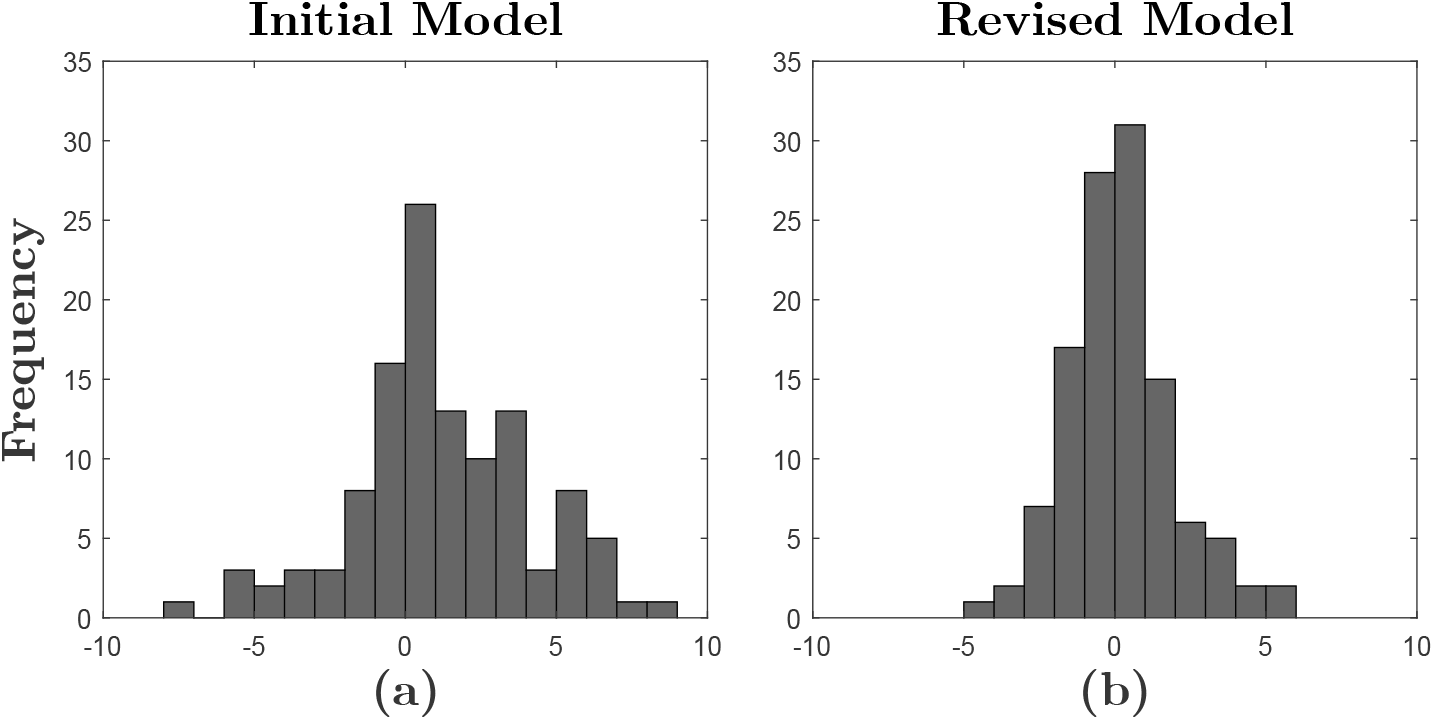
Residual distribution for both initial and revised models. The Y-axis means the times of the occurrence of residuals at a certain range. The residual is calculated as *E*^*j*^ = *A*^*j*^ *− M* (*t*^*j*^) (**a**) for the initial model, and (**b**) for the revised model.

Due to the unchanged parameters at Stage I, the calcium dynamics follow the same patterns as in the initial model. Figure 4 depicts the dynamics of Stages II and III for iTBS 600, imTBS600, and cTBS600 with prior contraction. The initial model demonstrates significant deviations (*RMSE*_Case1,Ini_ = 3.10, *SD*_Case1,Ini_ = 2.62) compared to the revised Huang model (*RMSE*_Case1,Rev_ = 1.73, *SD*_Case1,Rev_ = 1.63). Moreover, for imTBS600, the amount of each *substance* is approximately equal (Figure 4, first row, middle column), and its after-effect is not obvious as suggested in the previous experimental work (Figure 4, second row, middle column) (Huang et al., 2005).

**Figure 4:**
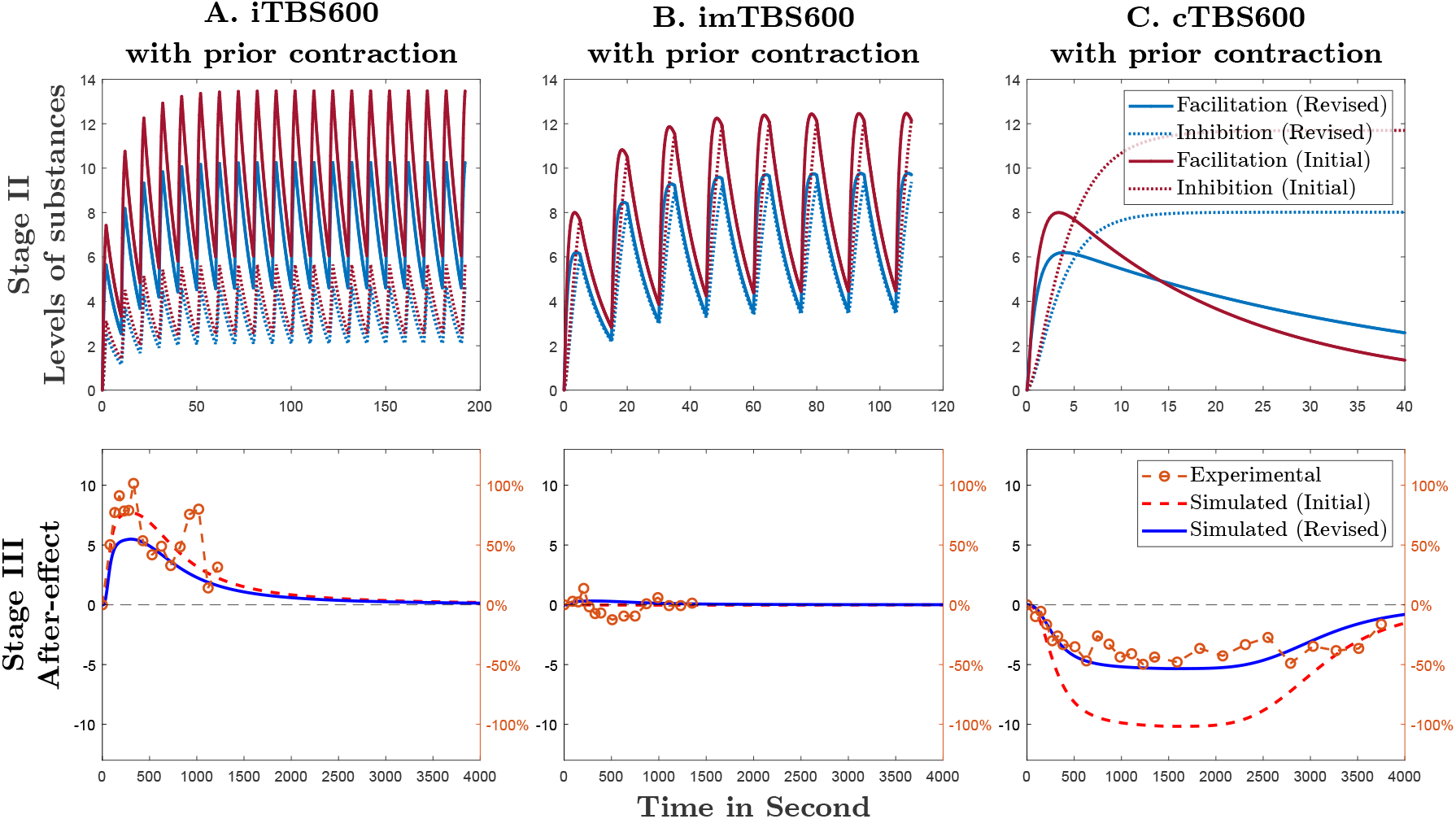
Case 1: Simulated results of the revised and the initial Huang model for **A**) iTBS600 (HU05), **B**) imTBS600 (HU05), and **C**) cTBS600 (HU05) with prior voluntary muscle contraction. Stage II shows the levels of variation of facilitatory and inhibitory *substances* during the TBS intervention. Stage III depicts the time course of the after-effects. These experimental measurements are extracted from the previous work by Huang et al. (2005). The left *Y* axis represents the simulated results in arbitrary units, while the right *Y* axis is the percentage of change of the MEP amplitudes for the experimental results.

Figure 5 illustrates the results of the revised model for cTBS300 with or without prior contraction and cTBS600 without prior contraction. Both polarities of these after-effects are well described by the revised model in comparison with the initial model. The revised model has a better performance (*RMSE*_Case2,Rev_ = 1.55, *SD*_Case2,Rev_ = 1.50) than the initial model (*RMSE*_Case2,Ini_ = 3.51, *SD*_Case2,Ini_ = 2.94) in terms of the after-effect magnitudes.

**Figure 5:**
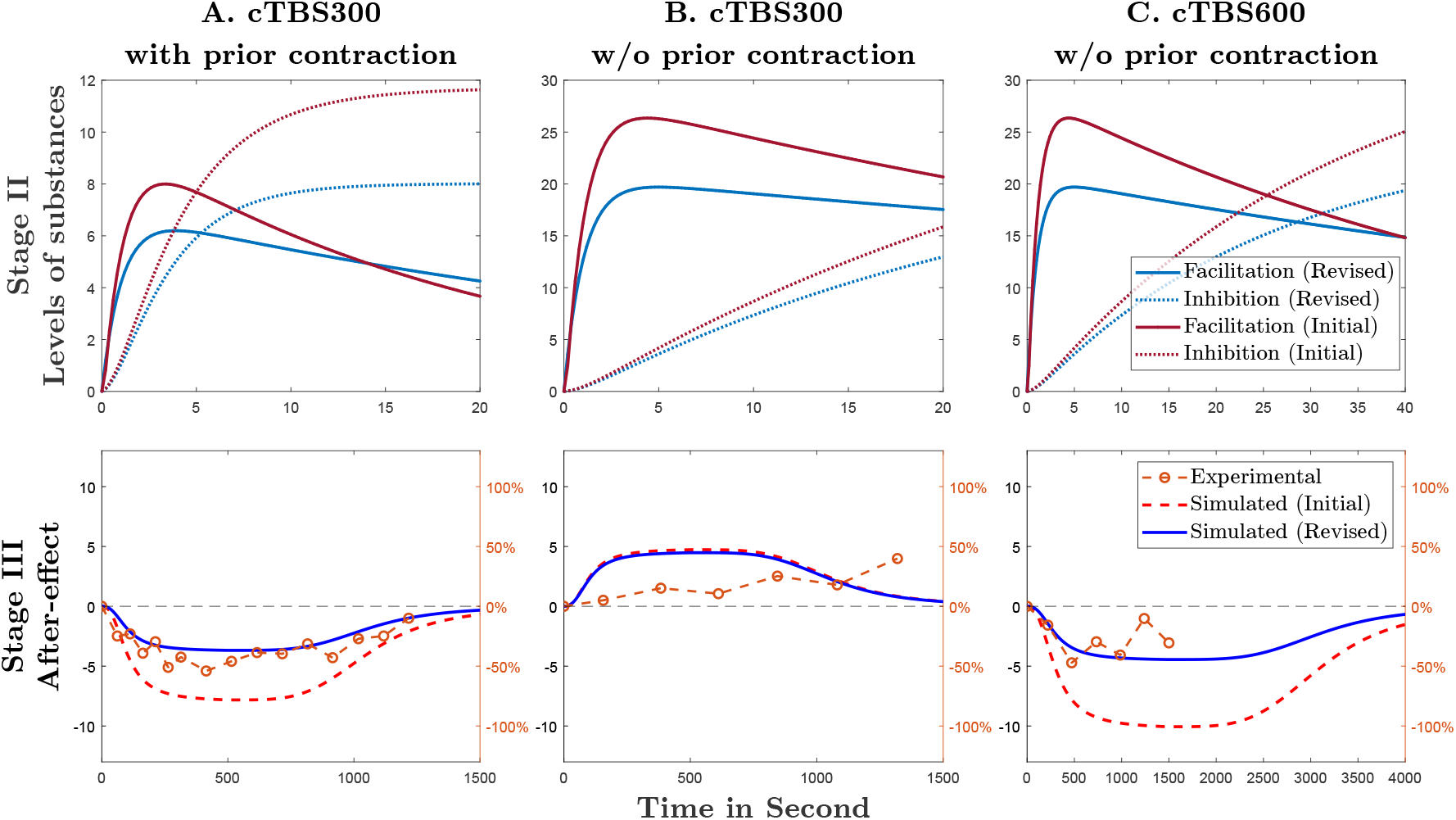
Case 2: Simulated results of the revised and the initial Huang model for **A**) cTBS300 with (HU05) or **B**) without (GE08) prior voluntary muscle contraction and **C**) cTBS600 (GE08) without prior contraction. Stage II shows the levels of variation of facilitatory and inhibitory *substances* during the TBS intervention. Stage III depicts the time course of the after-effects. These experimental measurements are extracted from the previous studies by Huang et al. (2005) and Gentner et al. (2008). The left *Y* axis represents the simulated results in arbitrary units, while the right *Y* axis is the percentage of change of the MEP amplitudes for the experimental results.

In addition, Figure 6 displays the after-effects of TBS immediately followed by voluntary contraction. Post-intervention muscle contraction reversed the after-effect of cTBS300, whereas it enhanced the after-effect of iTBS600 in experiments and simulations. The residuals for both models are within an acceptable range (*RMSE*_Case3,Ini_ = 2.96, *SD*_Case3,Ini_ = 2.91 and *RMSE*_Case3,Rev_ = 2.17, *SD*_Case3,Rev_ = 2.21).

**Figure 6:**
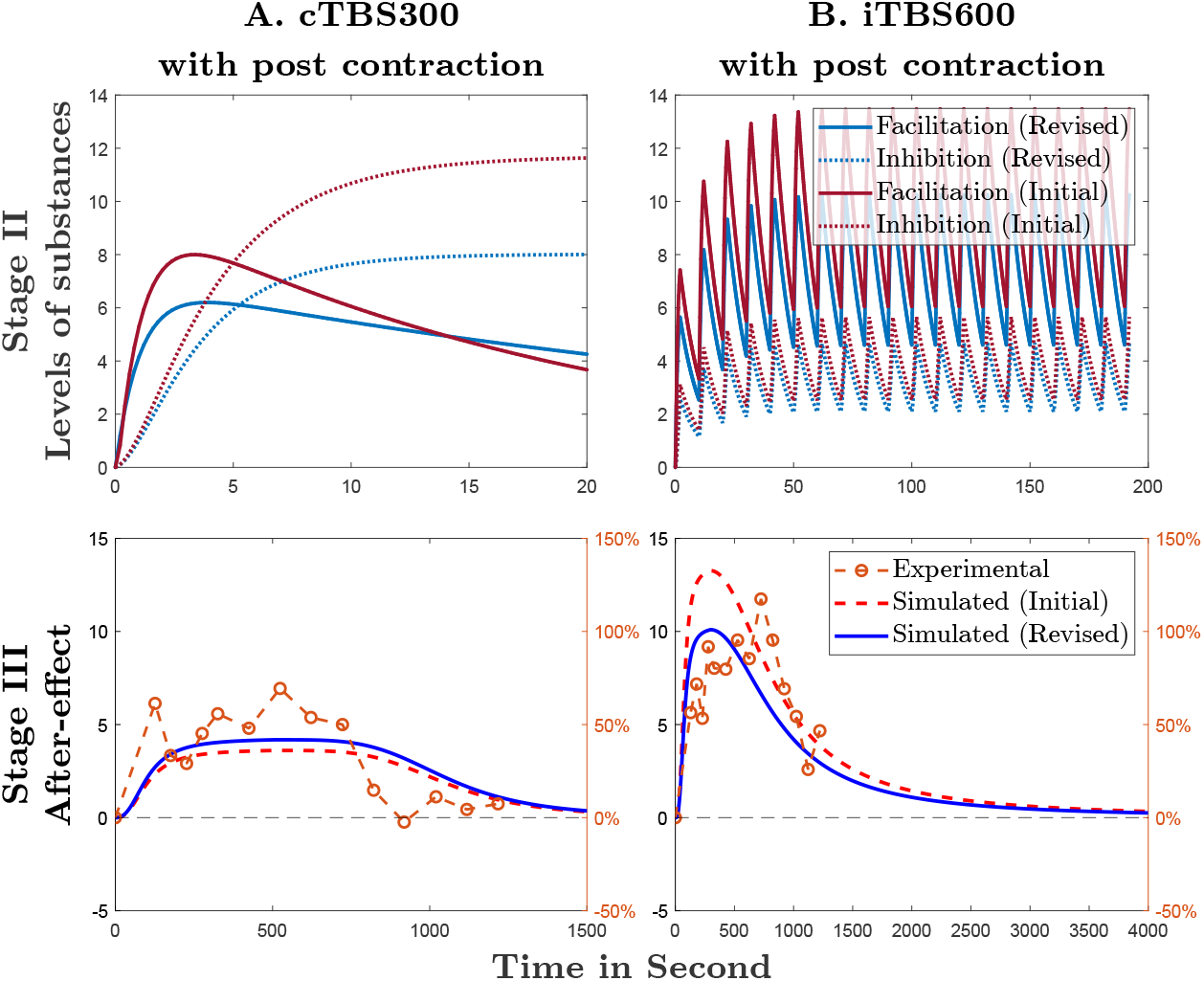
Case 3: Simulated results of the revised and the initial Huang model for **A**) cTBS300 (HU08) and **B**) iTBS600 (HU08) with prior as well as post contraction. Stage II shows the levels of variation of facilitatory and inhibitory *substances* during the TBS intervention. Stage III depicts the time course of the after-effects. These experimental measurements are extracted from the previous studies by Huang et al. (2008). The left *Y* axis represents the simulated results in arbitrary units, while the right *Y* axis is the percentage of change of the MEP amplitudes for the experimental results.

## 5. Discussion

### 5.1. Overall performance and limitations

We corrected as well as re-calibrated the Huang model and demonstrated its better ability to describe the after-effects of three theta-burst TMS protocols (specifically cTBS, imTBS, iTBS) under various experimental conditions. The revised Huang model supports the hypothesis that TBS simultaneously induces a mixture of inhibitory and facilitatory effects and further illustrates how TBS protocols affect the polarities and time courses of after-effects by inducing different [Ca^2+^]_*i*_ elevation patterns. However, this model is simple as it only incorporates the mechanisms and dynamics of glutamatergic synapses and postsynaptic Ca^2+^ influx through NMDA receptors.

The model only regards few protocol characteristics to describe the very specific protocol of TBS, specifically the number of pulses/bursts/trains and the inter-train/inter-stimulus intervals, naive and not defined for any other protocol changes. Further, as a first-order design with static parameters and fixed time-independent decay dynamics, neither the initial nor the revised model can at present correctly predict TBS-induced after-effects induced by stimulation over 600 pulses (Gentner et al., 2008; Gamboa et al., 2011; Doeltgen, Ridding, 2011; Haeckert et al., 2021).

### 5.2. Interpretation of the parameters

At Stage I, Ca^2+^ influx with prior contraction is significantly lower than that without prior contraction, which suggests that endogenous neural activation moderates neural responses to external stimulation. The decay constant *k* indicates the speed of Ca^2+^ removal from the cytoplasm by multiple ion pumps and exchangers (Berridge et al., 2000).

Due to the index error at Stage II, the initial computer code overestimates the rate of increase in [Ca^2+^]_*i*_, whose value is larger than that in the revised model. Hence, a corresponding increase of the value of *R*_f_ according to Table 2 for maintaining a sufficient increment of facilitatory *substances* after each burst in the calibration appears logic. The decay constant of facilitation *k*_f_ decreases with the correction, which means that the induced effects tend to remain longer than that in the initial model. In addition, the decay constant of inhibition *k*_i_ increases as does its proportionality constant *R*_i_. They indicate that the inhibition *substance* has a faster decline after bursts and a lower final steady level.

At Stage III, the summation of facilitatory and inhibitory *substances* could be interpreted by the dephosphorylation/phosphorylation of sites on the AMPA receptor subunit protein GluR_1_ (Lee et al., 2000; Castellani et al., 2001). The match of the initial model to the experimental data is worse than that of the revised one, which can be attributed to an improvement at Stage II of the revised model.

### 5.3. Further considerations

The majority of studies reported that rTMS can elicit suppression or enhancement of MEPs as an indirect way to access the excitability of the cortical circuits in the human motor cortex after stimulation (Carson et al., 2016). Although several lines of evidence indicate that TMS-induced after-effects and long-term changes of synaptic plasticity may share similar mechanisms (Huang et al., 2008; Wankerl et al., 2010; Di Lazzaro et al., 2008, 2005; Ziemann et al., 1996; Peinemann et al., 2000; Di Lazzaro et al., 2002, 2006; Liepert et al., 1997), it seems not plausible yet to name such effects LTP/LTD plasticity without direct evidence showing a strong link of such TMS-induced changes to synaptic plasticity (Cárdenas-Morales et al., 2010; Huang et al., 2017). Thus, we use the term *neuromodulation* to describe the alternation of neural activities induced by TMS.

Future mechanistic models of TMS neuromodulation in general should take into account a number of known mechanisms. Many studies of TBS-induced neuromodulatory effects have been performed *in vivo* and demonstrated that GABAergic mechanisms should be involved during TBS (Li et al., 2019). With the help of magnetic resonance spectroscopy (MRS), they reported that an inhibitory protocol (i.e., cTBS) elicits an increment of the concentration of GABA and has no significant impact on the concentration of glutamate/glutamine (Glx), whereas a facilitatory protocol (i.e., iTBS) can induce an obvious reduction of the GABA/Glx ratio (Stagg et al., 2009; Harrington, Hammond-Tooke, 2015; Iwabuchi et al., 2017; Kotak et al., 2017). Accordingly, these findings suggest that both GABAergic and glutamatergic neurotransmission are relevant for the afore-mentioned after-effects.

In addition to neurotransmitters, several studies demonstrated that the calcium influx though NMDA receptors may not be the only calcium source inducing after-effects. Grehl et al. (2015) recently suggested that patterned stimulation protocols (e.g., theta-burst) may mainly induce a Ca^2+^ release from intracellular stores (especially the endoplasmic reticulum) rather than Ca^2+^ influx from the extracellular space in neurons *in vitro*. Moreover, Wankerl et al. (2010) indicated that L-type voltage-gated calcium channels may contribute to the changes of [Ca^2+^]_*i*_ during TBS and act as molecular switches mediating metaplasticity induced by voluntary muscle contraction.

Further parameters of TBS pulses may become significant in experiments and also future models, such as the pulse strength, the pulse shape, the number of pulses per TBS, and the frequency of pulses in bursts (Nyffeler et al., 2008; Gamboa et al., 2010; Wu et al., 2012; Goldsworthy et al., 2012; Sasaki et al., 2018; Strzalkowski et al., 2019; Ozdemir et al., 2021; Goetz et al., 2016; D’Ostilio et al., 2016; Shirota et al., 2017; Li et al., 2022).

## 6. Conclusion

In this paper, we corrected inconsistencies between the computer code and the formulation in the text of Huang et al. as well as re-calibrated the index-adjusted Huang model through a non-convex optimisation to the same experimental data used for the initial model. The revised Huang model likewise supports the hypothesis that a mixture of facilitatory and inhibitory effects are simultaneously induced during TBS. Furthermore, it captures the pattern as well as time course of the relevant experimental results. Although the model design is highly specific and simple, it is transparent, well-structured, and provides both a quantitative model for several parameters of TBS and a basic model foundation for future refinement.

## 7. Dedication

This work is dedicated to Y.-Z. Huang, who pioneered theta-burst stimulation with TMS and introduced a very first model to reproduce as well as explain its effect but could not contribute any more to this work due to his unfortunate early passing.

## Appendix A. Supplementary Material

An implementation of the model in MATLAB accompanied by instructions for its use is available at https://github.com/BIOMAKE/Revised_Huang_Model.

## Footnotes

1 MATLAB code is attached to the supplement material of the initial publication and also used for the plots in the article Huang et al. (2011).

2 Plots from the given literature were digitized using *WebPlotDigitizer* by Ankit Rohatgi.

